# scaLR: a low-resource deep neural network-based platform for single cell analysis and biomarker discovery

**DOI:** 10.1101/2024.09.19.613226

**Authors:** Saiyam Jogani, Anand Santosh Pol, Mayur Prajapati, Amit Samal, Kriti Bhatia, Jayendra Parmar, Urvik Patel, Falak Shah, Nisarg Vyas, Saurabh Gupta

## Abstract

**Purpose:** Single-cell RNA sequencing (scRNA-seq) is producing vast amounts of individual cell profiling data. Analysis of such datasets presents a significant challenge in accurately annotating cell types and their associated biomarkers. scRNA-seq datasets analysis will help us understand diseases such as Alzheimer’s, Cancer, Diabetes, Coronavirus disease 2019 (COVID-19), Systemic Lupus Erythematosus (SLE), etc. Recently different pipelines based on machine learning (ML) and Deep Neural Network (DNN) methods have been employed to tackle these issues utilizing scRNA-seq datasets. These pipelines have arisen as a promising resource and are capable of extracting meaningful and concise features from noisy, diverse, and high-dimensional data to enhance annotations and subsequent analysis. Existing tools require high computational resources to execute large sample datasets.

**Methods:** We have developed a cutting-edge platform known as scaLR (Single Cell Analysis using Low Resource) that efficiently processes data in batches, and reduces the required resources for processing large datasets and running NN models. scaLR is equipped with data processing, feature extraction, training, evaluation, and downstream analysis. The data processing module consists of sample-wise & standard scaler normalization and splitting of data. Its novel feature extraction algorithm, first trains the model on a feature subset and stores feature importance for all the features in that subset. At the end of this process, top K features are selected based on their importance. The model is trained on top K features, its performance evaluation and associated downstream analysis provide significant biomarkers for different cell types and diseases/traits.

**Results:** To showcase the capabilities of scaLR, we utilized several scRNA-seq datasets of Peripheral Blood Mononuclear Cells (PBMCs), Alzheimer’s patients, and large datasets from human and mouse embryonic development. Our findings indicate that scaLR offers comparable prediction accuracy and requires less model training time and compute resources than existing Python-based pipelines and frameworks. Moreover, scaLR efficiently handles large sample datasets (*>*11.4 million cells) with minimal resource usage (29GB RAM, 12GB GPU, and 8 CPUs) while maintaining high prediction accuracy and being capable of ranking the biomarker association with specific cell types and diseases.

**Conclusion:** We present scaLR a Python-based [platform](https://github.com/infocusp/scaLR), engineered to utilize minimal computational resources while maintaining comparable execution times to existing frameworks. It is highly scalable and capable of efficiently handling datasets containing millions of cell samples and providing their classification and important biomarkers.

## 1 Introduction

Single-cell RNA sequencing (scRNA-Seq) is increasingly prevalent due to its ability to provide more comprehensive data compared to bulk RNA-seq. Moreover, accessibility to scRNA-seq data is expanding with the development and optimization of the technology by different companies, resulting in increased throughput and reduced costs [1]. scRNA-Seq data analysis facilitates the identification of cell types based on their gene expression profiles across different biological samples [2]. Accurate classification of cells into types, sub-types, and states is imperative for precise single-cell analysis, particularly in disease classification, evaluation of therapeutic strategies, biomarker identification, and subsequent analysis [3]. Cell types annotation is mainly performed using marker genes, employing correlation-based techniques, utilizing supervised classification [4] and recently, DNN-based models have been developed to perform the annotations [5–7].

Clustering is a commonly used method that annotates cell types utilizing manually curated marker genes obtained from literature to assign cell types to clusters generated through unsupervised learning [8]. Selection of marker genes relies on researchers’ prior knowledge, which can introduce biases and inaccuracies. Additionally, marker genes may not be readily available for all cell types of interest, especially for novel cell types that lack established marker gene sets. Moreover, many cell types are defined by a combination of genes rather than a single marker gene [9]. Without a robust method to incorporate expression data from multiple marker genes, ensuring consistent and precise annotation of cell types to each cluster becomes challenging, time-consuming, laborious, and less scalable for large datasets [8, 10–12]. Tools such as cellMeSH [13], CellAssign [14], scCATCH [15], SCINA [16], SCSA [17], scSorter [18] and scType [19] perform clustering first and then assign a cell type identity to each cluster.

The correlation-based methods such as Scmap [20], SingleR [21], scMatch [22], CHETAH [23], measure correlation of gene expression profiles between the query samples and reference dataset [8] and mainly affected by the batch effect of platforms and experiments [24]. Numerous batch-effect correction methods are available, but distinguishing genuine biological diversity from technical disparities remains challenging, thereby preserving crucial biological variations poses a challenge [25].

Annotation of cell types using supervised/semi-supervised classification and DNN methods adhere to the traditional paradigm in machine learning, which identifies patterns in gene expression profiles and transfers labels from labeled to unlabeled datasets [26]. These methods such as ACTINN [27],Celltypist [28], devCellPy [29], scBalance [30], scvi-tools [31], scdeepinsight [32], SingleCellNet also known as pySingleCellNet [33] and cellPLM [7] have gained popularity recently due to their resilience to data noise and variability, as well as their independence from artificially chosen marker genes. It has been noticed that ML and DNN-based tools necessitate significantly high computational resources(*>*128GB RAM,*>*12GB GPU memory, and *>*8 CPUs), to handle datasets comprising *>*0.12 million single-cell samples having approx 22.5K genes.

To address this challenge, we’ve developed scaLR, which tested in low resources VM (128GB RAM, 12GB GPU, and 8 CPUs) and a Laptop (32GB RAM and 8 CPUs). This platform can process millions of cells by dividing the dataset into batches and using different layers of DNN models for training. Furthermore, it also provides downstream analysis of genes elucidating their specific roles in determining cell type, trait, and disease.

## 2 Methods

### 2.1 scRNA-seq data selection and download

PBMCs constitute a crucial component of the immune system, pivotal in combating a spectrum of infections stemming from harmful pathogens. They serve as a fundamental tool for investigating the immune response, infectious ailments, cancer, and the development of vaccines [2, 34]. To design and compare the scaLR platform performance PBMCs-Bacterial Sepsis (PBMCs-BS) dataset was used. Furthermore, to showcase the efficacy of scaLR and compare it with other pipelines, we downloaded and curated various PBMC studies, COVID-19, human breast, and embryonic development scRNA-seq datasets. Along with it human brain scRNA-seq data of different cortexes is downloaded to identify the Alzheimer’s disease-specific biomarkers (Table 1). These individual studies were primarily sourced from Cellxgene [35] and the Single-cell portal of Broad Institute [25].

**Table 1.** List of scRNA-Seq datasets used to test the scaLR performance and compare with other frameworks.

| Datasets | Number of Cells | Number of Genes | Number of Cell-type | Reference |
| --- | --- | --- | --- | --- |
| PBMCs-BS<sup>1</sup> | 1,26,351 | 22,858 | 6 | [36] |
| PBMCs-Normal<sup>2</sup> | 6,85,024 | 36,771 | 25 | [37] |
| PBMC-C19-F<sup>3</sup> | 8,36,148 | 37,292 | 17 | [38] |
| PBMC-C19-H<sup>4</sup> | 9,00,360 | 45,821 | 35 | [39–41] |
| PBMC-AIDA<sup>5</sup> | 10,58,909 | 36,161 | 33 | [42] |
| PBMCs-SLE<sup>5</sup> | 12,63,676 | 30,867 | 11 | [43] |
| HBSFC<sup>7</sup> | 2,28,479 | 38,353 | 8 | [44, 45] |
| HBPFC<sup>8</sup> | 3,60,190 | 50,197 | 9 | [46–48] |
| HBCA<sup>9</sup> | 21,22,065 | 15,166 | 26 | [49] |
| HED<sup>10</sup> | 40,62,980 | 46,397 | 72 | [50] |
| MED<sup>11</sup> | 114,41,407 | 45,854 | 134 | [51] |

<sup>1</sup>PBMCs-BS: PBMCs-Bacterial Sepsis
<sup>2</sup>PBMCs-Normal: PBMCs from healthy human
<sup>3</sup>PBMC-C19-F:PBMCs-COVID-19-Flu (PBMCs of COVID-19, sepsis and flu patient)
<sup>4</sup>PBMC-C19-H:PBMCs-COVID-19-Harmonized(PBMCs from various COVID-19 studies were pooled together and harmonized)
<sup>5</sup>PBMCs-AIDA: PBMCs-Asian Immune Diversity Atlas
<sup>6</sup>PBMCs-SLE: PBMCs-Systemic lupus erythematosus
<sup>7</sup>HBSFC: Human brain superior frontal gyrus
<sup>8</sup>HBPFC: Human brain prefrontal cortex
<sup>9</sup>HBCA:Human breast cell atlas
<sup>10</sup>HED:Human embryonic development
<sup>11</sup>MED:Mouse embryonic development

### 2.2 The scaLR platform

Various ML and DNN-based automated frameworks have been recently developed for cell type classification, which requires a significant amount of GPU memory and RAM to process raw data and generate the model that classifies cells. To address this, we developed scaLR, a Python-based platform that provides cell annotations and associated important biomarkers. scaLR comprises data processing, feature extraction, training, and evaluation & downstream analysis (Figure 1).

**Fig. 1.**
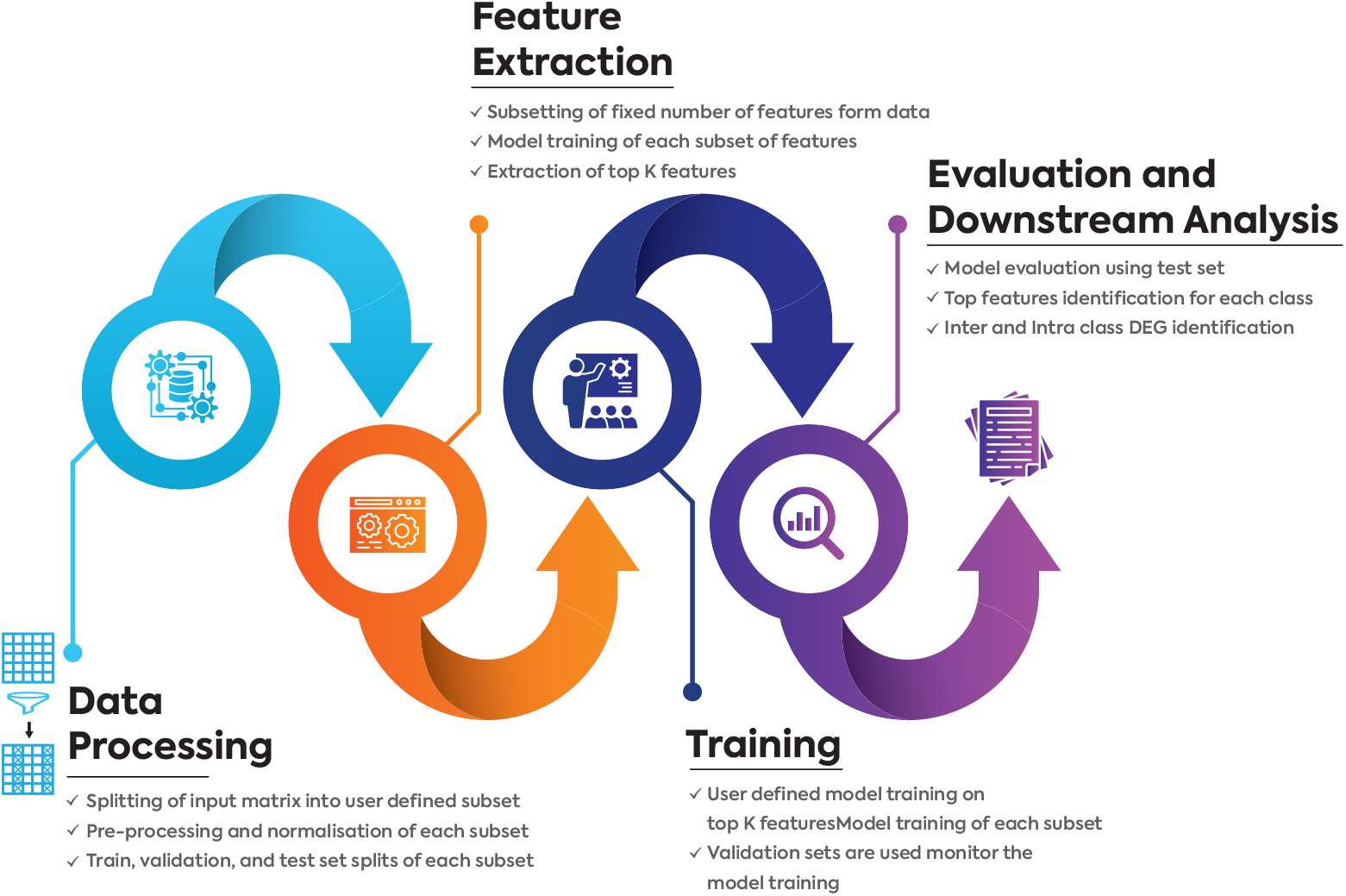
Schematic representation of different steps of scaLR.

#### 2.2.1 Data Processing

Large samples of the TPM matrix accompanied by metadata in the h5ad file are divided into subsets and subset-wise preprocessing (normalization) is adopted to optimize resource usage and storage. Subsequently, batch correction (if the user opts) is handled by the scaLR dataloader module on each subset to remove the noise from gene expression and eliminate batch effects [52]. This approach effectively manages RAM load as per system specifications, without impacting computational time. Each subset sample is divided into training, validation, and test sets for further platform steps. Notably, the subset sample size of the selected X features from all features matrix serves as a critical factor in memory consumption across the platform. This enables reducing the subset size significantly reducing memory usage without introducing excessive computational overhead.

#### 2.2.2 Feature Extraction

The normalized input dataset comprises thousands of sparse features, emphasizing the need for faster, more efficient, and superior model training. Identification of important features from each is an important section of the platform. To address this need, we have developed the unique **Feature Extraction Algorithm** 1. This algorithm partitions the input dataset into a user-defined number of feature subsets, and a single-layered neural network model is fitted for a given training dataset. Subsequently, the validation set is employed to assess the model’s effectiveness. The weights for each subset of features are then stored. This process iterates for multiple feature subsets, ensuring no repetition. Each subset is used to fit exactly one model, with each model trained on a distinct subset of features. The weights matrix generated from each model is utilized to evaluate the contribution of features toward predicting specific classes.

It is important to note that each model was trained in the same condition, using the same optimizer, and all their weights were initialized to zero, to ensure symmetry and fairness. All subset weight matrices combine to form a weight matrix consisting of weights for all features across all classes. Then, the mean of absolute weights of each feature across all classes represents the score of a feature contributing towards a prediction. The features are then ranked according to their scores (higher is better) calculated by the scorer call of the module. After top-k features are chosen to train the final model further explained in Figure 2.

**Fig. 2.**
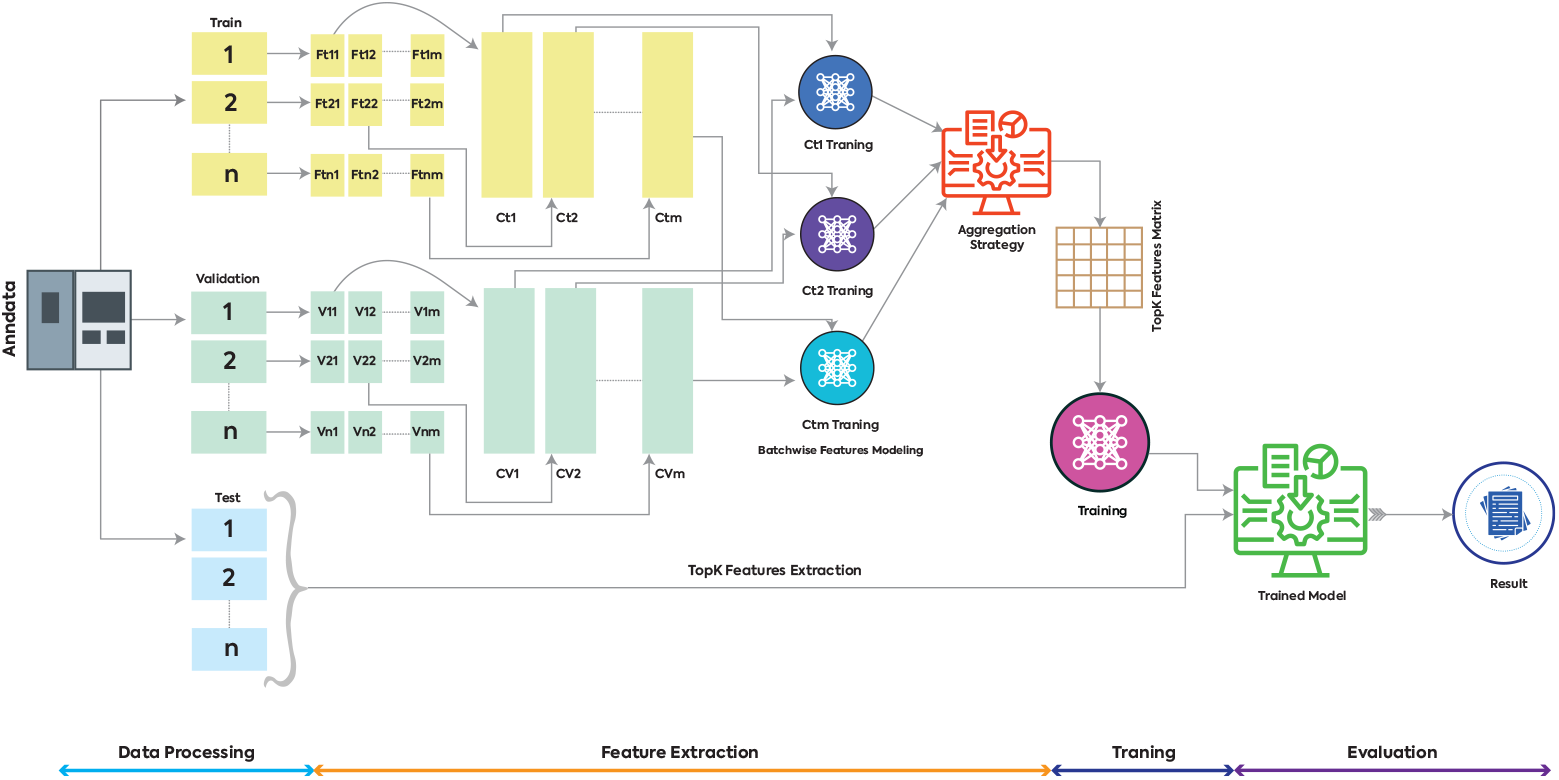
Overview of scaLR, a multilayered machine-learning platform for cell type classification and disease-specific features identification. Along with it performs downstream analysis of genes to relevant biomarkers. User-defined top-K features can be extracted using a single or multilayer layer neural network for all feature subsets by performing batch-wise data processing and feature extraction. These top-K features are used in final DNN model training and evaluation of these models is performed using a test set.

##### Algorithm 1

Feature Extraction Algorithm used in scaLR

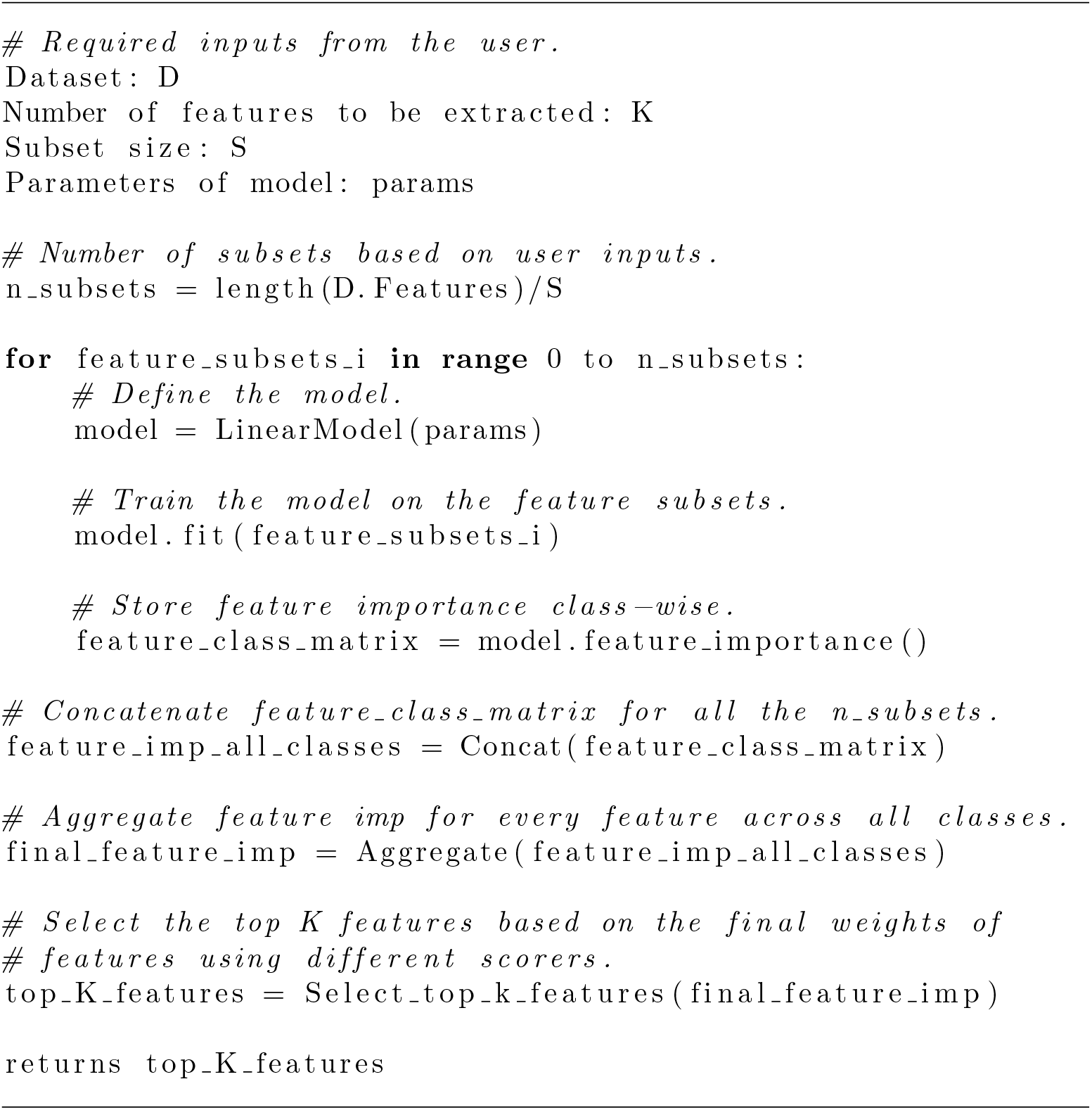

#### 2.2.3 Training

During DNN model training, the top K feature dataset is utilized to train the model, while the validation dataset is employed to monitor performance with early stopping to identify the optimal model. To adhere principle of minimal resource consumption, the training data is directly read from disk, and learning is conducted in batches, thereby ensuring efficient processing of considerably large datasets containing tens of thousands of features. This approach allows scaLR to be executed on any machine.

#### 2.2.4 Evaluation and Downstream Analysis

The trained model is then evaluated on the test set using metrics such as precision, recall, f1-score, and accuracy scores. A detailed classification report will show the model’s performance in each class. In the case of the linear model, model weights are used to identify the rank of top features (100 default) that contributed to predicting a particular class, while for multiple layered models, the SHapley Additive exPlanations (SHAP) algorithm [53] is used to identify class-specific top features. scaLR downstream analysis module embedded with classes/functions of gene recall curves, ROC & AUC curves, and heatmap generation. However, DGE analysis can be performed using Pseudobulk and Linear mixed-effects model (LMEM) approaches, and the module will generate their respective volcano plots.

### 2.3 scaLR performance comparison with different pipelines

To compare the performance of scaLR with other pipelines, we used both all features and the top 3,500 features extracted by the scaLR feature extraction module from the PBMCs-BS dataset (Table 1). This enabled us to evaluate the results obtained across different ML and DNN pipelines, which primarily include:

ACTINN employs NN with three hidden layers and is trained on datasets with predetermined cell types. It uses the trained parameters to predict cell types in other datasets [27].

CellTypist is an automated cell type annotation tool utilizing logistic regression classifiers optimized by the stochastic gradient descent algorithm. We have experimented with three variants of this tool i.e. 1) using logistic classifier 2) using SGD classifier and 3) using feature selection method followed by SGD classifier with all default parameters mentioned in classifier’s scikit-learn packages [28].

cellPLM is the first pre-trained large language transformer capable of encoding inter-cellular relationships, integrating spatially-resolved transcriptomic data, and utilizing a justified prior distribution. For model training on the PBMCs-BS dataset, we used 200 epochs for all features and 100 epochs for the top 3,000 selected features [7]. devCellPy offers various parameters that can be tuned based on the input dataset.

We used a variant that trains all layers without cross-validation and metrics, skipping the timepoint prediction. Additionally, initial parameter fine-tuning was turned off, allowing the tool to run with the standard default parameters set in the tool itself [29]. SingleCellNet or PySingleCellNet classifies cell types and states within heterogeneous cell populations by employing top-pair transformation followed by training a random forest classifier. It demonstrates the ability to classify across platforms and species with comparable sensitivity and specificity scores [33].

scvi-tools are designed for probabilistic modeling on scRNA-seq data and they utilize the inference procedure for scVI and scANVI models rely on NN, and stochastic optimization and employ variational autoencoders (VAEs) to infer latent representations of single-cell data in low-dimensional space [31].

scBalance employs a standard neural network architecture with 3-4 densely connected layers, incorporating batch normalization and dropout for better generalization, and uses the ELU activation function. Before training, we opted to scale the data and applied a weighted sampling technique to emphasize rare cell types [30].

In addition to comparing the cell annotation accuracy of scaLR with these tools, we also compare the top 100 identified genes of each cell type predicted by scaLR and cell Typist (all variant run) with cell-specific biomarkers identified by various sc-RNAseq-based studies and downloaded from CellMarker2.0 [54]. We can not extract the features or there is no explanation about how to get the top features for each cell type from generated models scBlance, scvi, devcellpy, and cellPLM.

To assess the performance and robustness of the scaLR platform, we have analyzed large sample datasets of different tissues of humans and mice. We also compared the DGE across various cell types in PBMCs-SLE and PBMC-C19-F datasets using the scaLR-enabled DEG module which uses the top 5,000 features provided by the model as well as using whole input data. scaLR is also capable of identifying disease-specific biomarkers using its novel feature extraction algorithm and associated model scorer for this we have used the human brain-superior frontal gyrus (HBSFC) and prefrontal cortex(HBPFC)(Table 1) datasets of normal and Alzheimer patients. To validate identified disease-specific top 10 genes we have performed literature mining of each gene for both cortex datasets.

## 3 Results

### 3.1 ScaLR

The scaLR platform consists of four primary modules those are data processing, feature extraction, model training, and evaluation & downstream analysis. The data processing module is designed to handle large sample datasets and segment them into training, testing, and validation sets. After that, it undergoes preprocessing of the dataset by performing sample-wise and/or standard scale normalization. The feature extraction modules play a pivotal role in scaLR’s uniqueness, as they identify class-specific top K features from each subset through single or multilayer DNN models. In the features selection model training phase, these feature subsets are trained into a DNN model. After all feature subsets are trained, the feature importance is evaluated across all features using Linear or SHAP scorer, and important features are selected. Then, the new model is trained on these top K important features and it undergoes evaluation using precision, recall, F1-score, and accuracy metrics alongside various reports like roc-auc [55] and gene recall curve on cell type and state annotations. Furthermore, essential biomarkers associated with distinct classes and differentially expressed genes (DEGs) between disease and normal cell types are identified. scaLR is implemented in Python, and comprehensive information about this platform can be seen and downloaded from our [github repo](https://github.com/infocusp/scaLR).

### 3.2 Performance evaluation of scaLR using all features

scaLR has a unique feature extraction capability, allowing users to select the top K features of the input dataset or process all features for classification using various models. The prediction accuracy of scaLR for cell type and state was compared with existing ML and DNN-based frameworks using all features of the PBMCs-BS dataset (Table 2). The results show that scaLR achieves similar accuracy to scVI-tools (scVI and scANVI) [31], SingleCellNet [33], CellTypist-Logistic, CellTypist-SGD, CellTypist-SGD+FS-TRP [28], ACTINN [27], and devCellPy [29] for cell type classification using a single-layer neural network. Additionally, scaLR performs better than scBalance [30], devCellPy [29], and cellPLM [7]. For cell state classification, scaLR shows the same accuracy as CellTypist-Logistic and cellPLM, and better accuracy than scBalance and SingleCellNet. ACTINN, CellTypist-SGD, CellTypist-SGD+FS-TRP, scVI-tools (scVI and scANVI), and devCellPy show comparatively better accuracy, possibly because we hyper-tuned the parameters of these frameworks to achieve the best accuracy, while in the case of scaLR, we did not optimize any of the parameters for all features run on PBMCs-BS dataset. It has been evident using all feature scVI-tools cannot execute due to its creating all features (22,858) latent spaces, which requires more than 128GB RAM and 12GB GPU. However, analysis was performed using the default number of latents (30). We gradually increased the number of latents up to 3,500 managing to stay within the available GPU memory. Similarly, cellPLM reduces the feature matrix to the top 3,000 features to run the models. In conclusion, the comparison indicates that scaLR performs better than existing NN-based frameworks in terms of accuracy when all features are used for cell-type annotation.

**Table 2.** Predicted accuracy of different pipelines on all features of PBMCs-BS dataset.

|  | Cell Type | Cell State |
| --- | --- | --- |
| Classifier | Accuracy | Accuracy |
| scaLR | 0.933 | 0.825 |
| scvi-tools(scVI & scANVI)<sup>1</sup> | 0.940 | 0.861 |
| SingleCellNet<sup>2</sup> | 0.939 | 0.812 |
| CellTypist-Logistic<sup>3</sup> | 0.940 | 0.821 |
| CellTypist-SGD<sup>4</sup> | 0.931 | 0.850 |
| CellTypist-SGD+FS-TRP<sup>5</sup> | 0.930 | 0.872 |
| scBalance<sup>6</sup> | 0.850 | 0.674 |
| devCellPy<sup>7</sup> | 0.940 | 0.852 |
| ACTINN<sup>8</sup> | 0.938 | 0.857 |
| CellPLM<sup>9</sup> | 0.889 | 0.821 |

<sup>1</sup>Used no of latent=30, no of layers=2, scVI epochs=100 and scANVI epochs=25
<sup>2</sup>Random forest using nTrees=500
<sup>3</sup>CellTypist using logistic classifier
<sup>4</sup>CellTypist using using SGD
<sup>5</sup>CellTypist using feature selection method followed by SGD classifier
<sup>6</sup>learning rate=0.005, minibatch size=128, epochs=50
<sup>7</sup>multi=softprob, eta: 0.2, max\_depth=6, subsample=0.5, colsample\_bytree=0.5, eval\_metric=merror and seed=840
<sup>8</sup>Weighted\_sampling=True
<sup>9</sup>NN model with top 3000 features run for 200 epochs

### 3.3 Comparison scaLR extracted top 3,500 features as input for different pipelines

In this comparison, we extracted the top 3,500 features of the PBMCs-BS dataset using the scaLR feature extraction algorithm that uses an SGD optimizer. We ran different pipelines using these features to evaluate prediction accuracy, memory use (maximum memory usage during the pipeline run), and wall clock time (end-to-end pipeline execution time) in a VM instance with 12GB GPU, 128GB RAM, and 8 CPUs. This experiment results indicate that wall clock time and maximum memory usage of scaLR for cell type and state are better than all other pipelines Table 3. However, prediction accuracy for cell type and state is better SingleCellNet [33], CellTypist(all variant run), scBalance [30], ACTINN [27], and cellPLM [7] and similar to scVI-tools (scVI and scANVI) [31], and devCellPy [29]. However, while running the devcellpy we have turned off the fine-tuning of parameters, probably leading to running the model faster using standard parameters defined inside the tool. Table 3 consists of each pipeline’s accuracy, time, maximum memory, and parameters used to execute these pipelines.

**Table 3.** Accuracy, wall clock time, and memory use in different pipelines executed on top 3,500 features with samples extracted from PBMC-BS dataset using scaLR features extraction module.

| Classifier | Cell Type (SGD) |  |  | Cell State (SGD) |  |  |
| --- | --- | --- | --- | --- | --- | --- |
|  | Accuracy | Time | Memory | Accuracy | Time | Memory |
| scaLR<sup>1</sup> | 0.941 | 2:33.15 | 1.70 | 0.840 | 6:23.39 | 1.70 |
| scvi-tools(scVI & scANVI)<sup>2</sup> | 0.940 | 21:25.53 | 3.76 | 0.860 | 22:47.22 | 3.77 |
| SingleCellNet<sup>3</sup> | 0.920 | 18:30.71 | 43.84 | 0.801 | 51:42.49 | 80.32 |
| scBalance | 0.790 | 14:12.31 | 6.35 | 0.690 | 14:16.51 | 6.35 |
| CellTypist-Logistic<sup>4</sup> | 0.930 | 51:45.10 | 4.38 | 0.830 | 2:30:31 | 4.38 |
| CellTypist-SGD<sup>5</sup> | 0.920 | 6:19.65 | 4.38 | 0.860 | 8:58.00 | 4.38 |
| CellTypist-SGD+FS-TRP<sup>6</sup> | 0.910 | 8:22.52 | 4.38 | 0.860 | 13:08.09 | 4.38 |
| devCellPy | 0.940 | 6:58.00 | 6.62 | 0.840 | 8:08.00 | 6.64 |
| ACTINN | 0.930 | 8:40.00 | 8.77 | 0.830 | 8:05.00 | 8.77 |
| CellPLM<sup>7</sup> | 0.890 | 9:05:19 | 3.86 | 0.840 | 7:02:41 | 3.74 |

\*SGD in cell type and state column indicate the input features.
**Time** Wall clock time in hours, minutes, seconds, and milliseconds (HH: MM: SS or MM: SS.MS) referred as end to end program run time.
**Memory** Maximum memory(gigabyte) utilized at any point of time.
<sup>1</sup>Used single layer, epochs=100
<sup>2</sup>Used number of latent=30, number of layers=2, scVI epochs=100 and scANVI epochs=25
<sup>3</sup>Random forest using nTrees=500
<sup>4</sup>CellTypist using logistic classifier.
<sup>5</sup>CellTypist using SGD.
<sup>6</sup>CellTypist using feature selection method followed by SGD classifier.
<sup>7</sup>CellPLM-The time is not included as we have run it on CPU and the model expects the GPU architecture for faster runs.

### 3.4 Evaluating the performance and robustness of scaLR on large sample datasets

To evaluate the performance and robustness of scaLR on large datasets, we selected a range of datasets with diverse features (approximately 22k to 46k) and sample sizes from 0.12 to 11.4 million cells (Table 1). scaLR has no constraints on the number of data samples that can be used for training, as we perform feature subsetting for feature extraction (if opted) and batch-wise data training. This allows users to train a model based on their available resources. Our goal is to assess the capability of other tools like CellTypist, devCellPy, SingleCellNet, scBalance, ACTINN, and CellPLM. We ran these pipelines on datasets with varying sample sizes (Table 1) using the same VM used in scaLR. Initially, we tested these tools on the PBMC-SLE dataset, which has over 1.2 million samples, using approximately 0.8 million samples for model training. Running each tool individually on this dataset caused memory crashes and failures to produce results. We then tested these frameworks on the PBMCs-Normal dataset, which has over 0.6 million cells, using approximately 0.4 million cells for model training. We observed that only CellPLM can run in this dataset while other tools require more than 128GB of memory, rendering them unable to function with limited or moderate resources. This issue arises because these tools attempt to load the entire dataset into memory simultaneously, which is not feasible for large datasets with limited resources.

On the other hand, scaLR enables the analysis of cross-cohort datasets exceeding 11.5 million cells, effectively managing batch effects generated by various single-cell sequencing platforms. These datasets include PBMC, human breast, and human and mouse embryonic development stages, each divided into the train, test, and validation sets as shown in Table 1. This experiment is crucial for ensuring the generalizability and reliability of scaLR in low-resource environments. Performance evaluation involves measuring the platform’s accuracy, precision, recall, F1-score (Supplementary Table 1-7), memory usage, and system runtime across diverse datasets, encompassing various biological conditions and technical variations (Table 4). Robustness evaluation assesses the scaLR’s ability to maintain consistent performance despite changes in input data characteristics, such as varying noise levels, batch effects, and data sparsity across all selected datasets. By systematically testing this platform on a comprehensive collection of datasets including benchmark datasets, one can identify strengths and limitations, ensuring that the platform is not overfitted to specific conditions. The associated results of these datasets are listed in supplementary tables and figures. This evaluation process is essential for confirming that the platform can accurately annotate cell types and subtypes, perform downstream analysis, and produce reliable results across different experimental settings, ultimately validating its applicability in various scRNA-seq studies.

**Table 4.** Performance evaluation and robustness check of scaLR in large sample datasets.

| Datasets | Training<sup>1</sup> |  |  | Evaluation<sup>2</sup> |  |  |
| --- | --- | --- | --- | --- | --- | --- |
|  | Samples size | Epoch | Layers | Accuracy | Memory | Time |
| PBMC-C19-F | 567761 | 23 | 5000,200,17 | 0.932 | 8.053 | 02:44:57 |
| PBMCs-Normal | 456682 | 40 | 5000,200,25 | 0.905 | 8.335 | 03:46:29 |
| PBMC-C19-H | 635132 | 15 | 5000,200,35 | 0.862 | 12.206 | 05:02:59 |
| PBMC-AIDA | 709215 | 28 | 5000,200,33 | 0.908 | 8.424 | 04:51:17 |
| PBMCs-SLE | 844488 | 30 | 5000,200,11 | 0.979 | 9.448 | 04:29:14 |
| HBCA | 1412546 | 17 | 5000,200,26 | 0.894 | 9.717 | 05:49:04 |
| HED | 3213236 | 26 | 5000,200,72 | 0.944 | 11.39 | 16:48:12 |
| MED | 8613510 | 10 | 5000,1000,300,134 | 0.927 | 29.072 | 63:27:59 |

**Note1** The training section covers factors like input sample size, epochs, and DNN layers, which ensure the effective execution of scaLR across various datasets. The evaluation section provides performance insights by presenting metrics such as accuracy, memory utilization, and total pipeline execution time.
**Note2** The performance of scaLR, including memory usage and overall runtime, may vary depending on the columns in the input AnnData object and the specified input parameters.
**Samples size** Number of samples in training.
**Epoch** Number of epochs used to converse the model out of 200 input epochs.
**Layers** Used DNN layers for the final model run.
**Time** Wall clock time in hours, minutes, seconds, and milliseconds (HH: MM: SS or MM: SS.MS) referred as end to end pipeline run time excluding downstream analysis time.
**Memory** Maximum memory(gigabyte) utilized at any point of time.

### 3.5 Features of scaLR

The scaLR is equipped with a range of advanced features designed to streamline and enhance the analysis of scRNA-seq data. It begins with the splitting of input datasets into the train, validation, and test subsets. Then user-defined normalization can correct the differences in sequencing depth and technical noise, ensuring consistent and comparable gene expression levels across cells. Feature extraction is the soul of this platform, enabling to loading of large sample datasets in different subsets and extracting the top K features using SGD optimizer. The platform uses these top-K features to accurately classify cells into distinct types and subtypes using NN models. It also provides cell type and disease-associated top 100 biomarkers. Integrated DGE analysis, allowing users to get DEGs between different conditions or cell types based on top K features of the model or in all features. To evaluate the performance of the model classification, ROC & AUC curve are used to illustrate the classification accuracy, plotting true positive rates against false positive rates, while gene recall curves assess the platform’s ability to recover known gene sets, providing insights into its sensitivity and robustness. Figure 3 shows the various downstream analysis plots generated by scaLR.Together, these features ensure a comprehensive, accurate, and reliable analysis of scRNA-seq data, making the platform an invaluable tool for researchers.

**Fig. 3.**
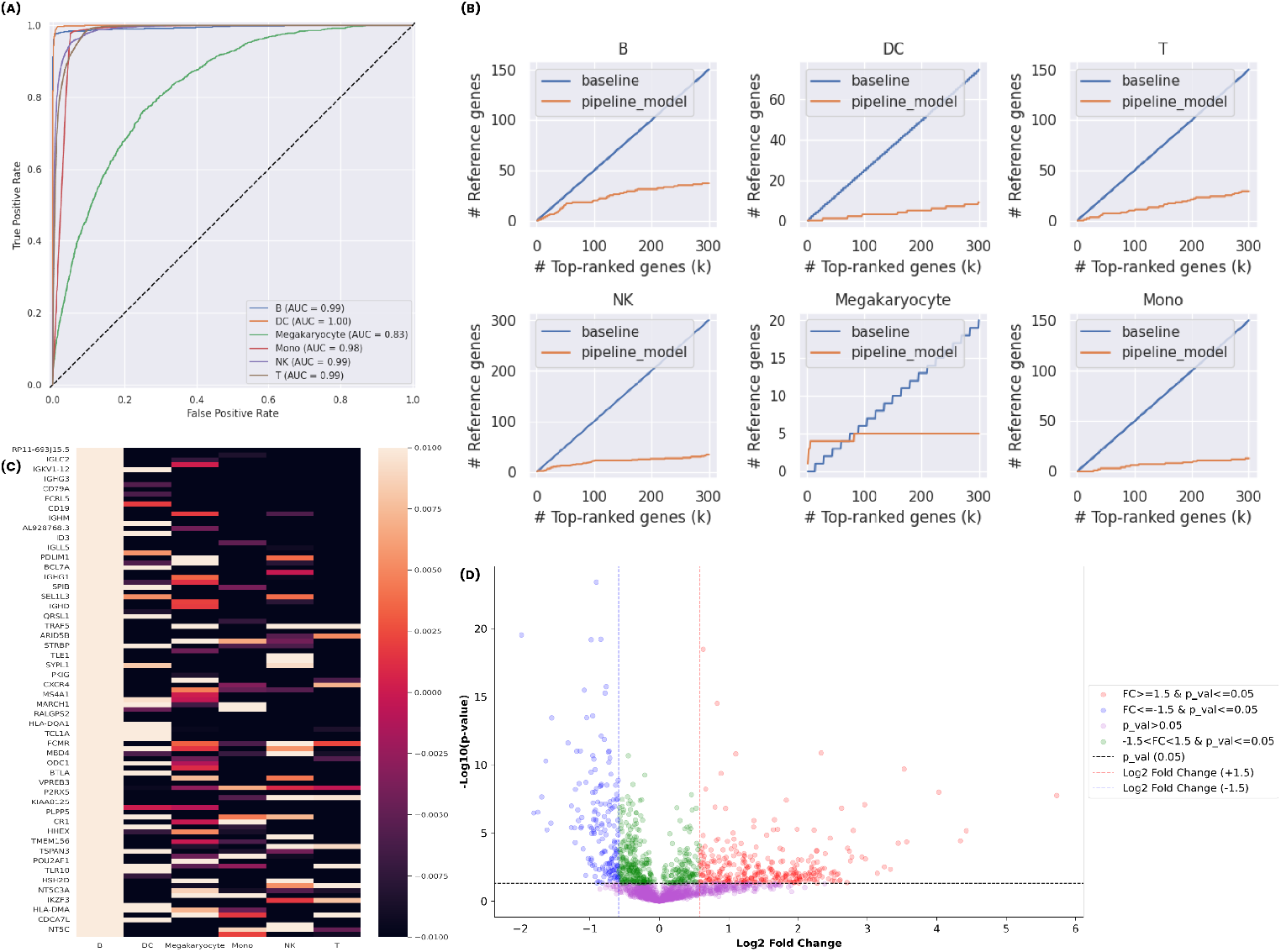
Different downstream analysis plots produced by scaLR (a) ROC & AUC curve showing the classification accuracy of predicted class, (b) Gene recall curves of cell-type specific biomarkers identified in top-ranked genes predicted by model w.r.t literature/reference genes, (c) An example heatmap indicates the top-ranked genes of a cell-type with their association in other cell type and (d) DGE plot showing differentially expressed genes (DEGs) for particular disease and normal condition

### 3.6 Celltype-specific biomarker discovery

Our comparative analysis of the top 100 cell type-specific biomarkers identified by scaLR and CellTypist-Logistic (the model with the highest accuracy) revealed significant overlaps and unique contributions from each tool in terms of literature or validated cell-specific biomarkers (Supplementary Table 8). Out of 223 reported T cell biomarkers, scaLR identified 21, while CellTypist predicted only 19. For megakaryocyte cells out of 39 biomarkers, scaLR predicted 10 while CellTypist’s only 5. Similarly, for monocytes, DC, and NK cells, scaLR predicted more biomarkers than CellTypist (Supplementary Figure 1). Only for B cells, CellTypist captures more biomarkers (33) compared to scaLR (32) within the top 100 features. DC showed the highest overlap between these tools across different cell types, while T and megakaryocyte cells had less overlap. We also plotted the gene recall for cell-specific biomarkers (Supplementary Table 8) as a reference concerning the identified biomarkers in the top 5,000 features of scaLR and CellTypist (Supplementary Figure 2). Both comparisons indicate that scaLR identifies more biomarkers than CellTypist. scaLR also generates a heatmap of the top 20 biomarkers identified through SHAP analysis for each cell type with their expression in other cells (Supplementary Figure 3).

Leveraging the power of SHAP, scaLR identifies the top 100 genes specific to each cell type, which are then compared to reference biomarkers for each cell type (Supplementary Table 8) using a gene recall curve for the PBMC-SLE (Supplementary Figure 4 and Supplementary Table 9) and PBMC-C19-F (Supplementary Figure 5 and Supplementary Table 10) datasets. These gene recall curves reveal that most biomarkers appear within the top 100 features for each cell type. In comparison to control patient samples, the scaLR DGE module pinpoints cell type-specific differentially expressed genes in PBMC-SLE (Supplementary Table 11 and Supplementary Figure 6) and PBMC-C19-F (Supplementary Table 12 and Supplementary Figure 7). This demonstrates scaLR’s accuracy in detecting cell-specific biomarkers, while other pipelines struggled to process these datasets on low-resource instances.

### 3.7 Diseases specific Biomarker Discovery

scaLR identifies the top 100 biomarkers for the HBSFG and HBPFC datasets, specific to Alzheimer’s disease and control groups, as listed in Supplementary Tables 13 and 14, respectively. Functional analysis of the top 10 genes reveals that scaLR effectively detects biomarkers related to Alzheimer’s disease and neurological disorders in both datasets. For the HBSFG dataset, seven of the top 10 genes are linked to neurological disorders, whereas in the HBPFC region dataset, only two of the top 10 genes are associated with such conditions. The functions of these genes are detailed in Supplementary Tables 13 and 14. While we annotated the functions of the top 10 genes, other highly ranked genes may also serve as potential biomarkers after further functional annotation.

## 4 Discussion

Recent advancements in single-cell sequencing techniques have significantly increased the generation of millions of single-cell datasets [1]. A necessary data analysis step is to perform cell-type annotation [2, 3], and associated biomarker discovery. With the publication of more comprehensive cell atlases, the popularity of auto-annotation tools has surged using different ML and NN models [26]. Despite this, existing frameworks face challenges in loading *>*0.5 million human single cells RNA-Seq data and performing the model training using all features in a low-resource compute system (12GB GPU memory, 128GB RAM, and 8 CPUs) with scalability, and compatibility. Our work introduces scaLR, a cutting-edge platform incorporating Sample batching, Normalization & Batch processing, Feature extraction, Feature ranking using different scores, and Gene recall modules. These modules enable scaLR to efficiently utilize minimal resources to train the model and perform different downstream analysis. Through a series of experiments with diverse scRNA-seq datasets varying in size, generation protocols, and levels of imbalance, we have demonstrated that scaLR is a low-resource deep neural network platform that excels in both annotation accuracy and execution speed with other existing frameworks.

Notably, we have compared scaLR performance of most of the widely used Python-based cell-type annotation tools such as scVI-tools (scVI and scANVI) [31], SingleCellNet [33], CellTypist [28], scBalance [30], ACTINN [27], devCellPy [29], scaLR has shown excellence in cell type annotation and features identification ability using all features and top 3,500 features (Table 2 and 3). In addition, in assessing the performance and robustness of scaLR in large sample datasets (Table 4), several key metrics were considered, including accuracy, memory usage, and end-to-end execution time. These metrics were evaluated across varying dataset sizes to gauge scaLR’s scalability and efficiency in handling high-dimensional data. Robustness checks were conducted by running the pipeline on multiple large datasets to ensure consistent results under different input conditions. Additionally, sensitivity analyses were performed by altering key input parameters, such as epochs and DNN layers, to evaluate the stability of the model’s predictions. This evaluation demonstrates that scaLR maintains high accuracy and manageable resource utilization, even with increased sample sizes, confirming its robustness for large-scale data analysis.

scaLR can also identify disease-specific biomarkers using control and disease-patient single-cell datasets (Supplementary Tables 10-14). Our analysis of the human brain’s two regions (SFG and PFC) datasets provides the top 100 ranked genes and most of them are possibly associated with Alzheimer’s disease and different neurological disorders (explained in the result section). We have performed functional annotation of the top ten ranked genes which are mainly in the HBSFG dataset Ferritin heavy chain 1 (FTH1) is involved in iron storage and regulation of oxidative stress. Its abnormal expression is linked to neurodegenerative diseases like Alzheimer’s and Parkinson’s due to iron accumulation [56]. Gene semaphorin-3C(SEMA3C) is involved in the emerging development of axons and dendrites of cortex [57, 58]. Diazepam binding inhibitor (DBI) is involved in the altered expression linked to anxiety, depression, and possibly epilepsy due to its role in neurosteroid production [59]. Overexpression stanniocalcin-1 (STC1) alleviates oxidative stress-induced injury, reduces neuroinflammation, and improves cognitive function [60]. Angiopoietin-like protein 4(ANGPTL4) plays a role in the process of white matter damage and cognitive impairment (CI) in patients with cerebral small vessel disease (CSVD)[61]. Histidine triad nucleotide-binding protein 1(HINT1) is involved in transcription regulation and apoptosis. Mutations in this gene are linked to neuropathy, schizophrenia, and cognitive impairments [62].

In the case of the HBPFC dataset, most of the top 10 ranked genes are long noncoding RNAs with unknown functions, while only two genes i.e. FERM RhoGEF and pleckstrin domain-containing protein1 (FARP-1) and Ribosomal protein S6 (RSP6) are found as a potential biomarker. FARP-1 is involved in Autism spectrum disorder (ASD) and cognitive impairment, particularly in its role in neural development [63]. RSP6 actively participates in protein synthesis and cell growth. Its phosphorylation is commonly used as a marker for neuronal activity [64].

In summary, we believe that scaLR is a valuable addition to cell class annotation and biomarker identification platform, particularly suited for low-resource environments and capable of efficiently handling datasets with millions of cells while providing accurate cell type annotations. Additional platform features include DGE analysis, gene recall curves, and heat maps of commonly associated cell-type genes. Future work could enhance its scalability and explore additional features to maintain its leading edge in scRNA-seq data analysis.

## Supplementary Information

### Supplementary Tables

**Supplementary Table 1**. Classification accuracy report for different cell types available PBMC-Normal dataset.

**Supplementary Table 2**. Classification accuracy report for different cell types available PBMC-SLE dataset.

**Supplementary Table 3**. Classification accuracy report for different cell types available PBMC-C19-H: PBMCs-COVID-19-Harmonized dataset.

**Supplementary Table 4**. Classification accuracy report for different cell types of PBMC-C19-F: PBMCs-COVID-19-Flu dataset.

**Supplementary Table 5**. Classification accuracy report for different cell types of HBCA dataset.

**Supplementary Table 6**. Classification accuracy report for different cell types of HED dataset.

**Supplementary Table 7**. Classification accuracy report for different cell types of MED dataset.

**Supplementary Table 8**. List of unique biomarkers for different cell types present in PBMC-BS, PBMC-SLE, and PBMC-COVID19-Flu datasets collected from CellMarker 2.0 (Download date 25 July 2024).

**Supplementary Table 9**. Top 100 cell type specific genes identified in PBMCs-COVID-19-Flu dataset using scaLR.

**Supplementary Table 10**. Top 100 cell type-specific genes identified in PBMCs-SLE dataset using scaLR.

**Supplementary Table 11**. Up- and down-regulated genes for each cell type concerning control samples in the PBMCs-SLE dataset.

**Supplementary Table 12**. Up- and down-regulated genes for each cell type concerning control samples in the PBMCs-COVID-19-Flu dataset.

**Supplementary Table 13**. Top 100 Alzheimer’s disease-specific genes identified in HBSFG dataset using scaLR-SHAP. Along with functional annotation of the top 10 genes of SHAP positive ranking for disease samples.

**Supplementary Table 14**. Top 100 Alzheimer’s disease-specific genes identified in HBPFG dataset using scaLR-SHAP. Along with functional annotation of the top 10 genes of SHAP positive ranking for disease samples.

### Supplementary Figures

**Supplementary Figure 1**. Venn Diagrams of (A) B cell, (B) T cell, (C) Monocytes, (D) Megakaryocytes, (E) DC and (F) NK cell of common and unique identified by scaLR and CellTypist biomarkers from the total biomarker for each cell type.

**Supplementary Figure 2**. Gene recall curve of identified biomarkers in top 5000 features concerning reference biomarker of cell types by scaLR and CellTypist.

**Supplementary Figure 3**. Heatmaps of top 20 genes identified by SHAP algorithms for (A) B cell, (B) T cell, (C) Monocytes, (D) Megakaryocytes, (E) DC and (F) NK cell and their respective expression in other cells identified by scaLR.

**Supplementary Figure 4**. Gene recall curve of identified biomarkers concerning reference biomarker of each cell identified by scaLR in top 5000 features of PBMC-SLE dataset.

**Supplementary Figure 5**. Gene recall curve of identified biomarkers concerning reference biomarker of each cell identified by scaLR in top 5000 features of PBMC-C19-F dataset.

**Supplementary Figure 6**. Volcano plots of differentially expressed genes (DEG) identified in different cell types concerning control samples in the PBMC-SLE dataset produced by the scaLR differential gene expression (DGE) analysis module.

**Supplementary Figure 7**. Volcano plots of differentially expressed genes (DEG) identified in different cell types concerning control samples in the PBMC-C19-F dataset produced by the scaLR differential gene expression (DGE) analysis module.

## Authors Contribution

This work was conceived and designed by: SG, AP, and NV; Entire platform development and deployment is performed by AP SJ, MP, SG, FS, and JP; Data analysis using this scaLR is done by AS, MP, AP, KB and SG Manuscript prepared by: SG, AP, FS, UP, and NV.

## Acknowledgements

We extend our appreciation to the IT Department for Inffocusp Innovations for providing the necessary resources and environment conducive to our research.

