## Supplementary Figure7 for "scaLR: a low-resource deep neural network-based platform for single cell analysis and biomarker discovery"

### Slide 1
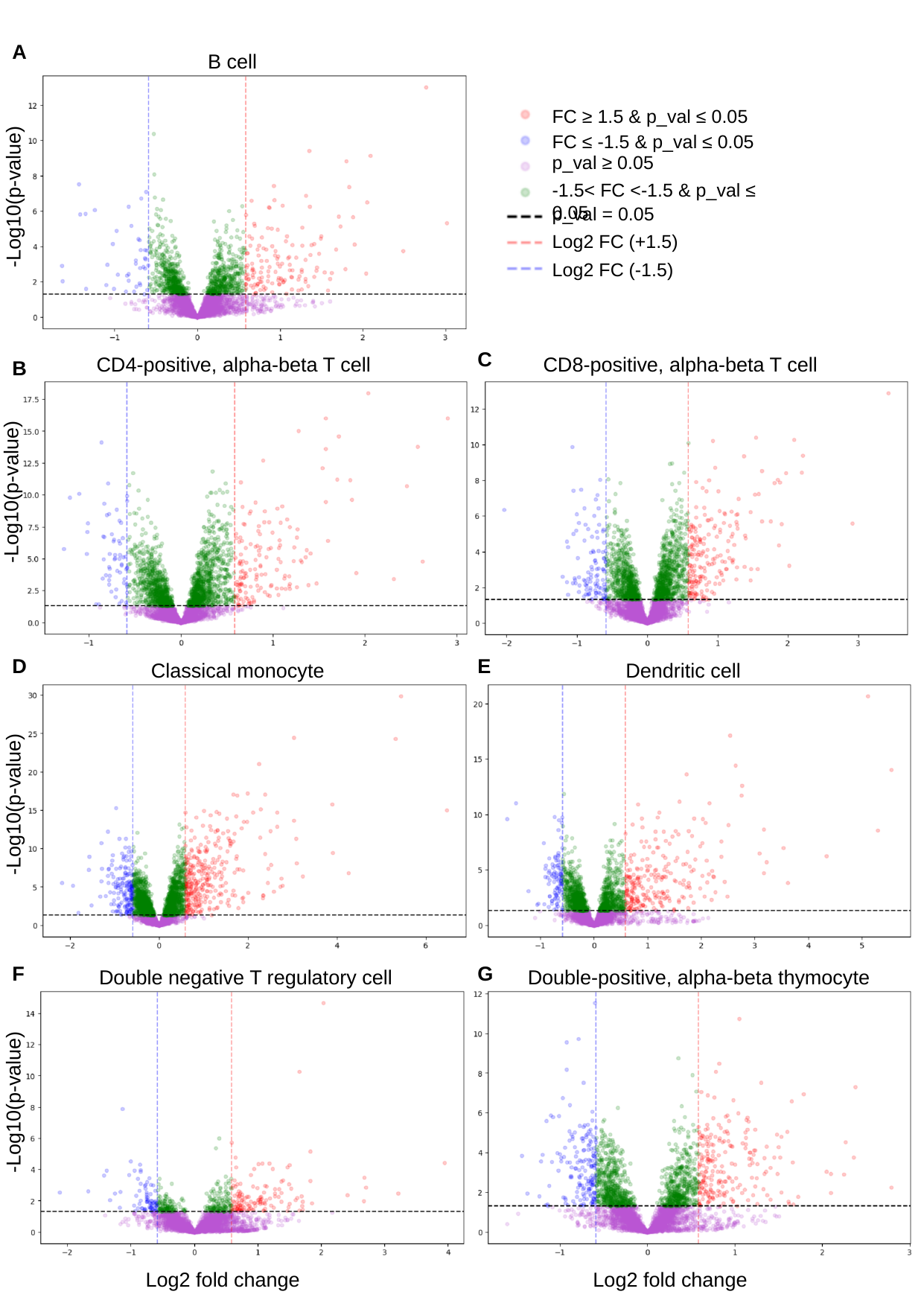

A
B cell
FC ≥ 1.5 & p_val ≤ 0.05
-Log10(p-value)
FC ≤ -1.5 & p_val ≤ 0.05
p_val ≥ 0.05
-1.5< FC <-1.5 & p_val ≤ 0.05
p_val = 0.05
Log2 FC (+1.5)
Log2 FC (-1.5)
C
CD4-positive, alpha-beta T cell
CD8-positive, alpha-beta T cell
B
-Log10(p-value)
D
E
Classical monocyte
Dendritic cell
-Log10(p-value)
F
G
Double negative T regulatory cell
Double-positive, alpha-beta thymocyte
-Log10(p-value)
Log2 fold change
Log2 fold change

### Slide 2
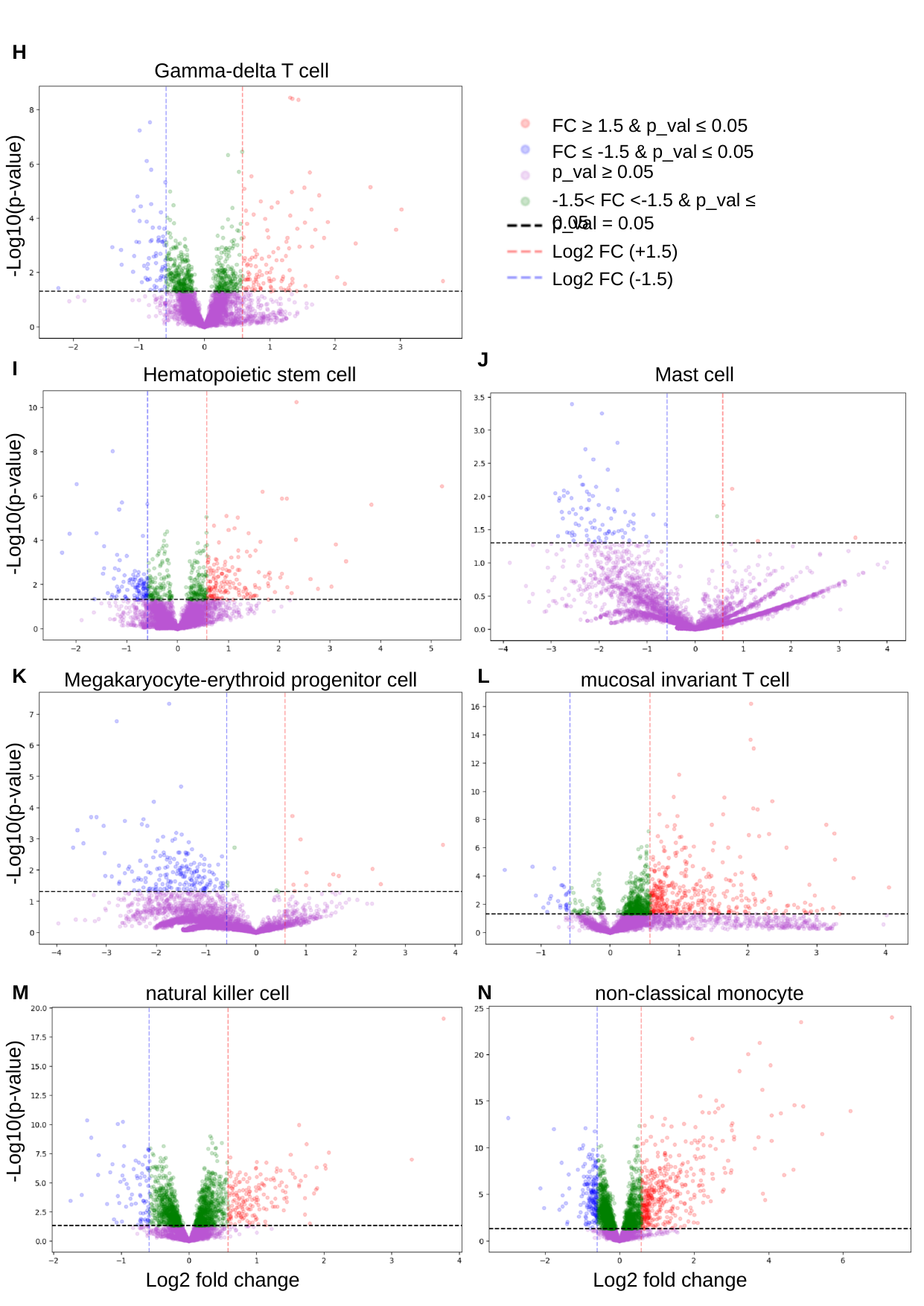

H
Gamma-delta T cell
FC ≥ 1.5 & p_val ≤ 0.05
-Log10(p-value)
FC ≤ -1.5 & p_val ≤ 0.05
p_val ≥ 0.05
-1.5< FC <-1.5 & p_val ≤ 0.05
p_val = 0.05
Log2 FC (+1.5)
Log2 FC (-1.5)
J
I
Hematopoietic stem cell
Mast cell
-Log10(p-value)
K
L
Megakaryocyte-erythroid progenitor cell
mucosal invariant T cell
-Log10(p-value)
M
N
non-classical monocyte
natural killer cell
-Log10(p-value)
Log2 fold change
Log2 fold change

### Slide 3
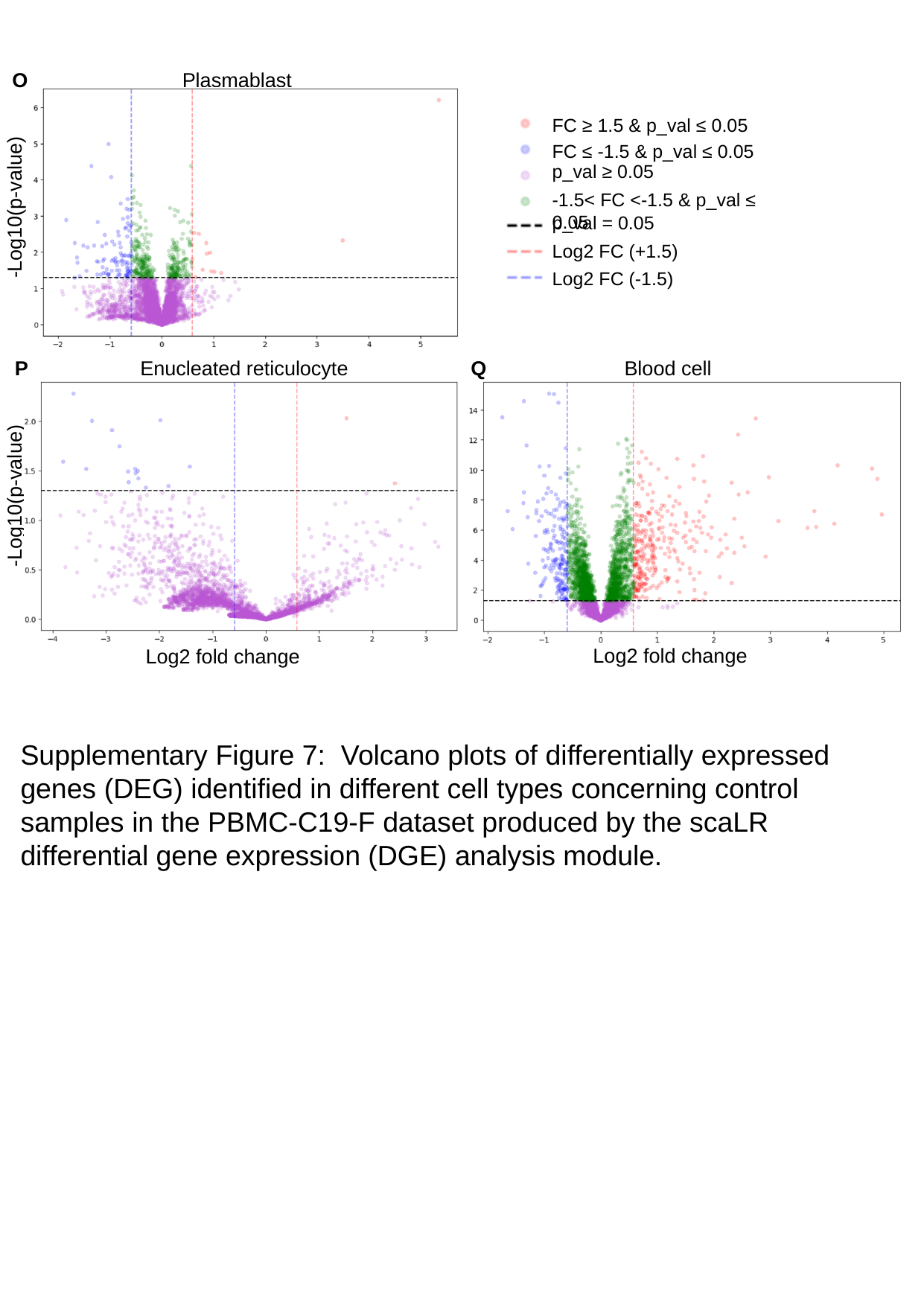

O
Plasmablast
FC ≥ 1.5 & p_val ≤ 0.05
-Log10(p-value)
FC ≤ -1.5 & p_val ≤ 0.05
p_val ≥ 0.05
-1.5< FC <-1.5 & p_val ≤ 0.05
p_val = 0.05
Log2 FC (+1.5)
Log2 FC (-1.5)
Enucleated reticulocyte
Blood cell
P
Q
-Log10(p-value)
Log2 fold change
Log2 fold change
Supplementary Figure 7: Volcano plots of differentially expressed genes (DEG) identified in different cell types concerning control samples in the PBMC-C19-F dataset produced by the scaLR differential gene expression (DGE) analysis module.
