## Supplementary Figure6 for "scaLR: a low-resource deep neural network-based platform for single cell analysis and biomarker discovery"

### Slide 1
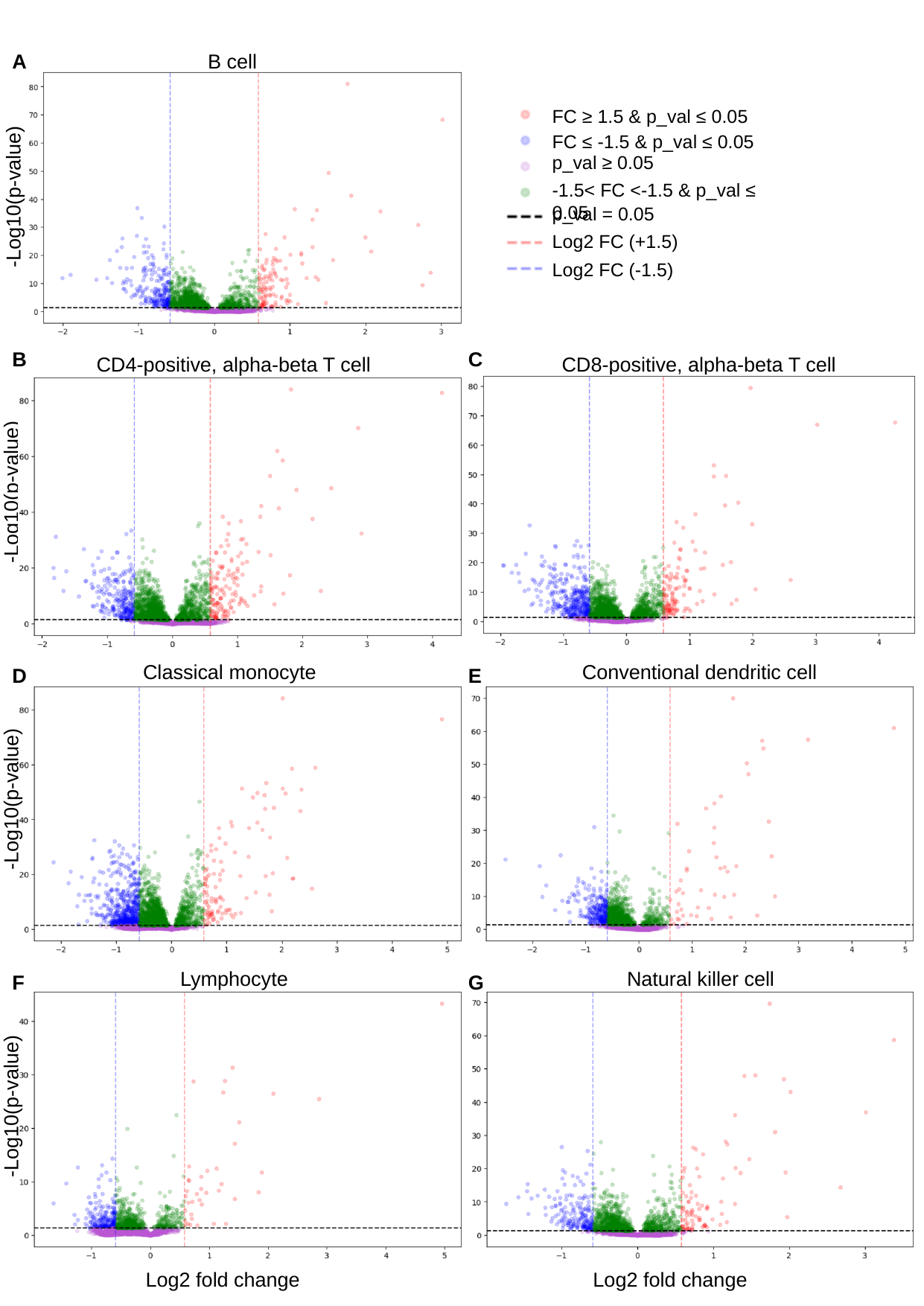

A
B cell
FC ≥ 1.5 & p_val ≤ 0.05
-Log10(p-value)
FC ≤ -1.5 & p_val ≤ 0.05
p_val ≥ 0.05
-1.5< FC <-1.5 & p_val ≤ 0.05
p_val = 0.05
Log2 FC (+1.5)
Log2 FC (-1.5)
B
C
CD4-positive, alpha-beta T cell
CD8-positive, alpha-beta T cell
-Log10(p-value)
Classical monocyte
Conventional dendritic cell
D
E
-Log10(p-value)
Lymphocyte
Natural killer cell
F
G
-Log10(p-value)
Log2 fold change
Log2 fold change

### Slide 2
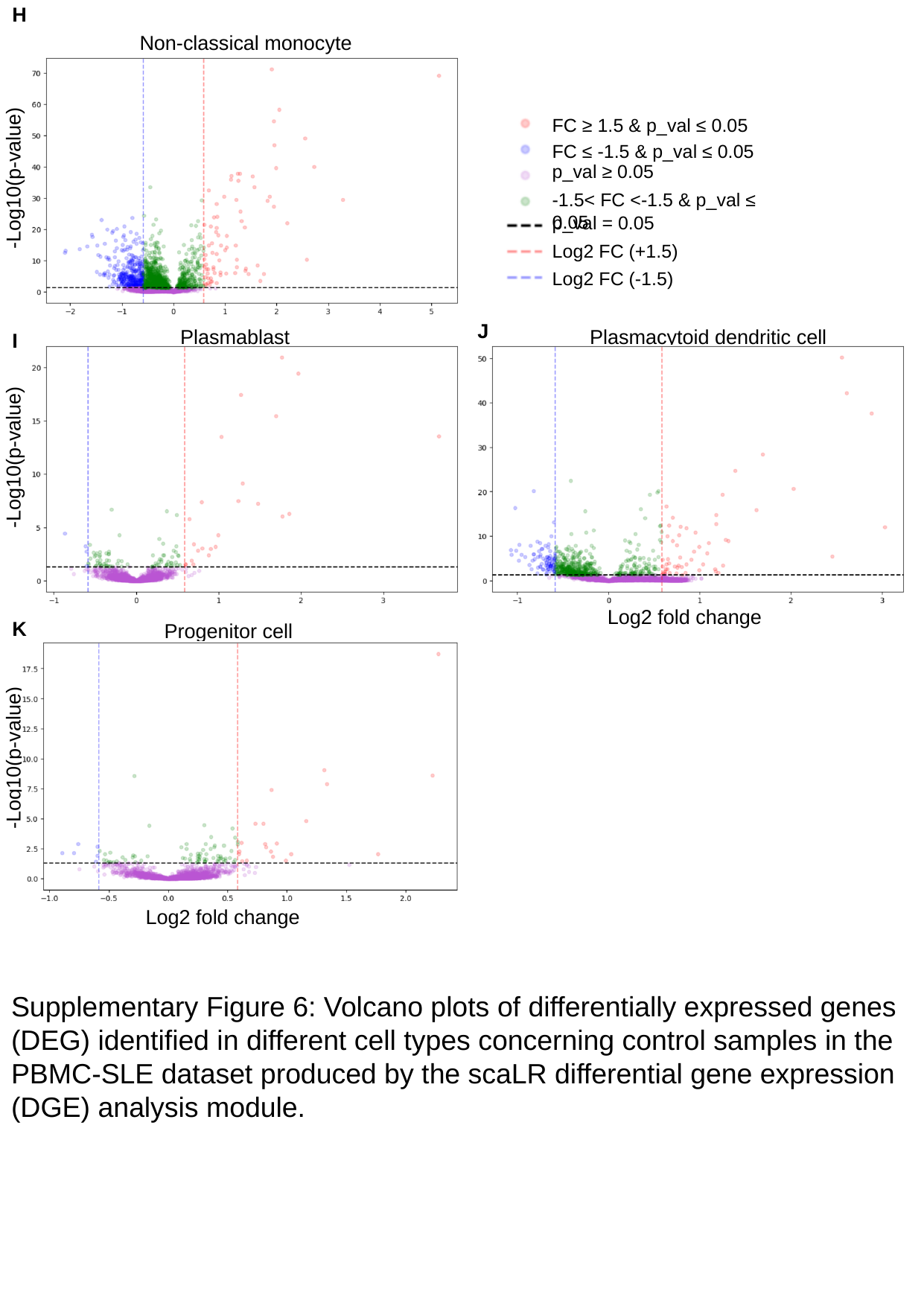

H
Non-classical monocyte
-Log10(p-value)
FC ≥ 1.5 & p_val ≤ 0.05
FC ≤ -1.5 & p_val ≤ 0.05
p_val ≥ 0.05
-1.5< FC <-1.5 & p_val ≤ 0.05
p_val = 0.05
Log2 FC (+1.5)
Log2 FC (-1.5)
J
Plasmablast
Plasmacytoid dendritic cell
I
-Log10(p-value)
Log2 fold change
K
Progenitor cell
-Log10(p-value)
Log2 fold change
Supplementary Figure 6: Volcano plots of differentially expressed genes (DEG) identified in different cell types concerning control samples in the PBMC-SLE dataset produced by the scaLR differential gene expression (DGE) analysis module.
