## Supplementary Figure5 for "scaLR: a low-resource deep neural network-based platform for single cell analysis and biomarker discovery"

### Slide 1
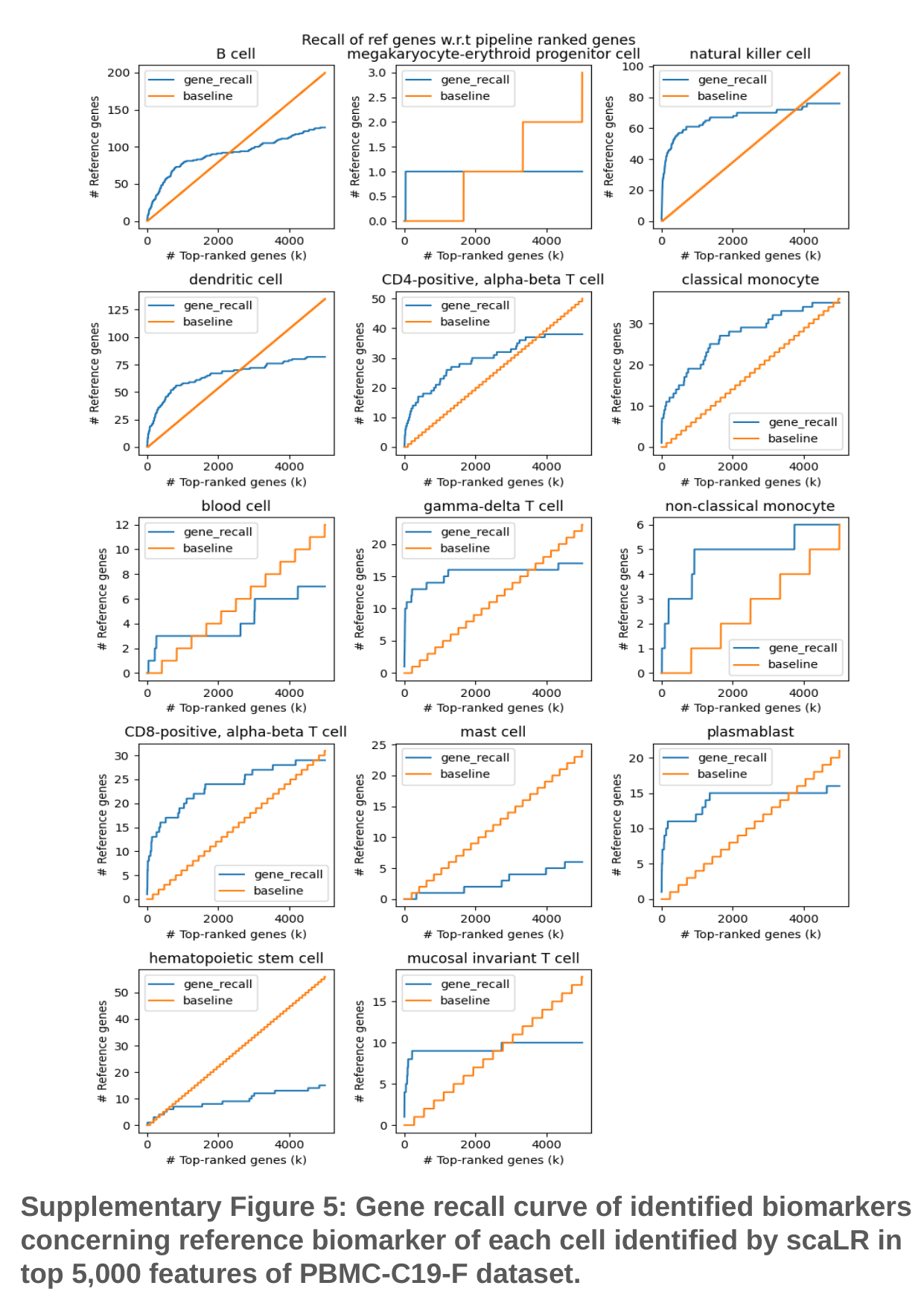

Supplementary Figure 5: Gene recall curve of identified biomarkers concerning reference biomarker of each cell identified by scaLR in top 5,000 features of PBMC-C19-F dataset.
