## Supplementary Figure2 for "scaLR: a low-resource deep neural network-based platform for single cell analysis and biomarker discovery"

### Slide 1
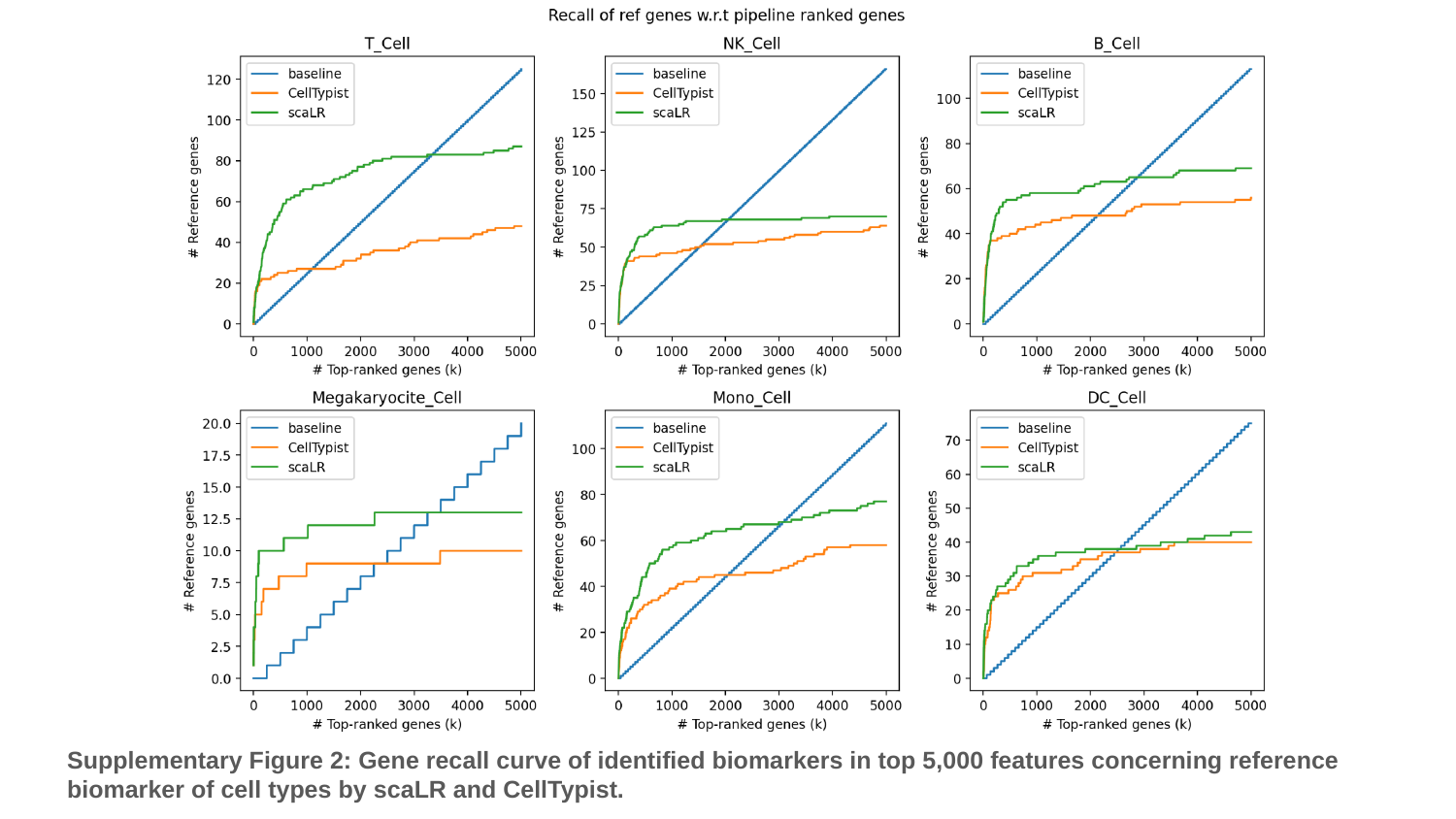

Supplementary Figure 2: Gene recall curve of identified biomarkers in top 5,000 features concerning reference biomarker of cell types by scaLR and CellTypist.
