## Supplementary Figure3 for "scaLR: a low-resource deep neural network-based platform for single cell analysis and biomarker discovery"

### Slide 1
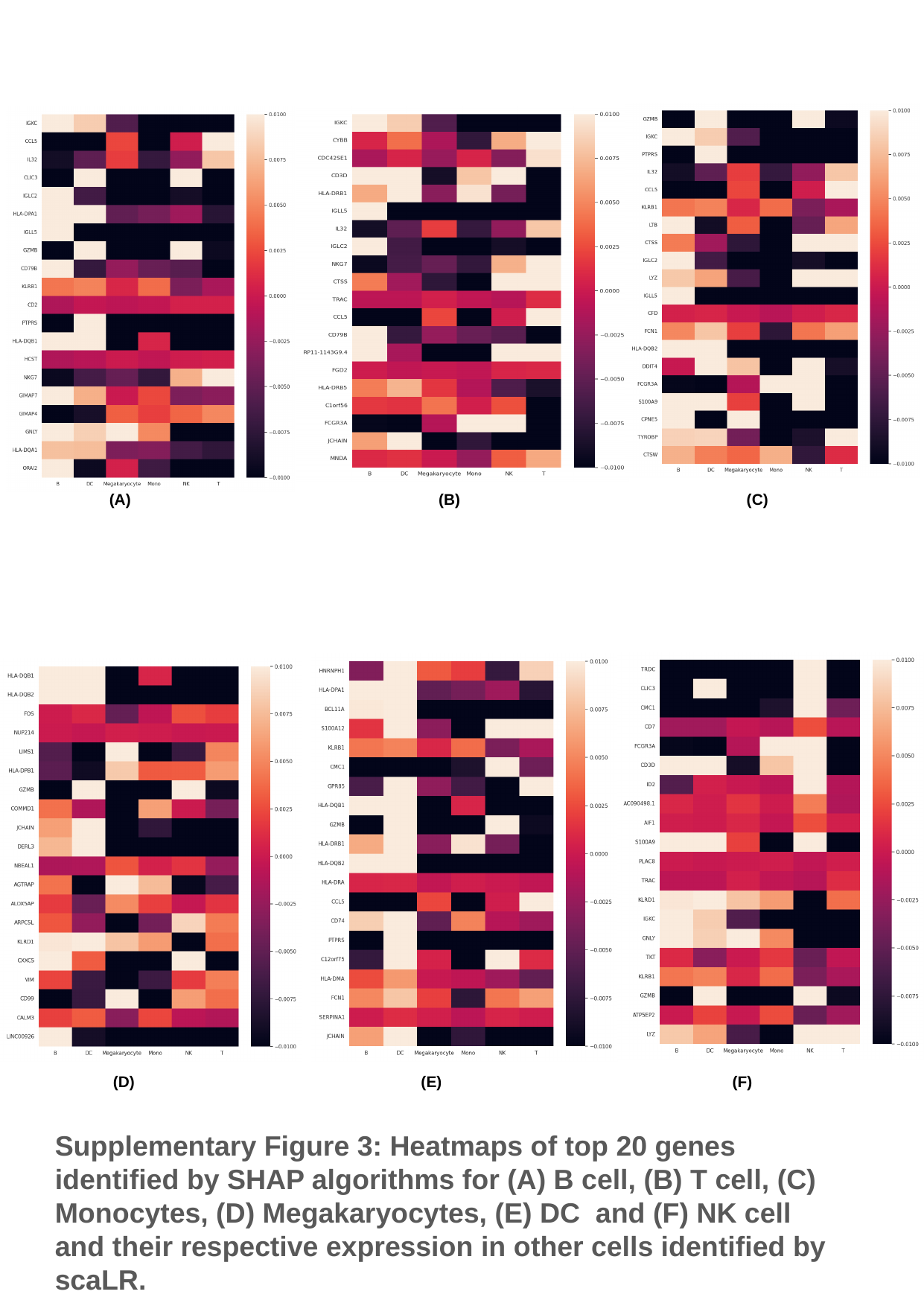

(A) (B) (C)
(D) (E) (F)
Supplementary Figure 3: Heatmaps of top 20 genes identified by SHAP algorithms for (A) B cell, (B) T cell, (C) Monocytes, (D) Megakaryocytes, (E) DC and (F) NK cell and their respective expression in other cells identified by scaLR.
