## Supplementary Figure1 for "scaLR: a low-resource deep neural network-based platform for single cell analysis and biomarker discovery"

### Slide 1
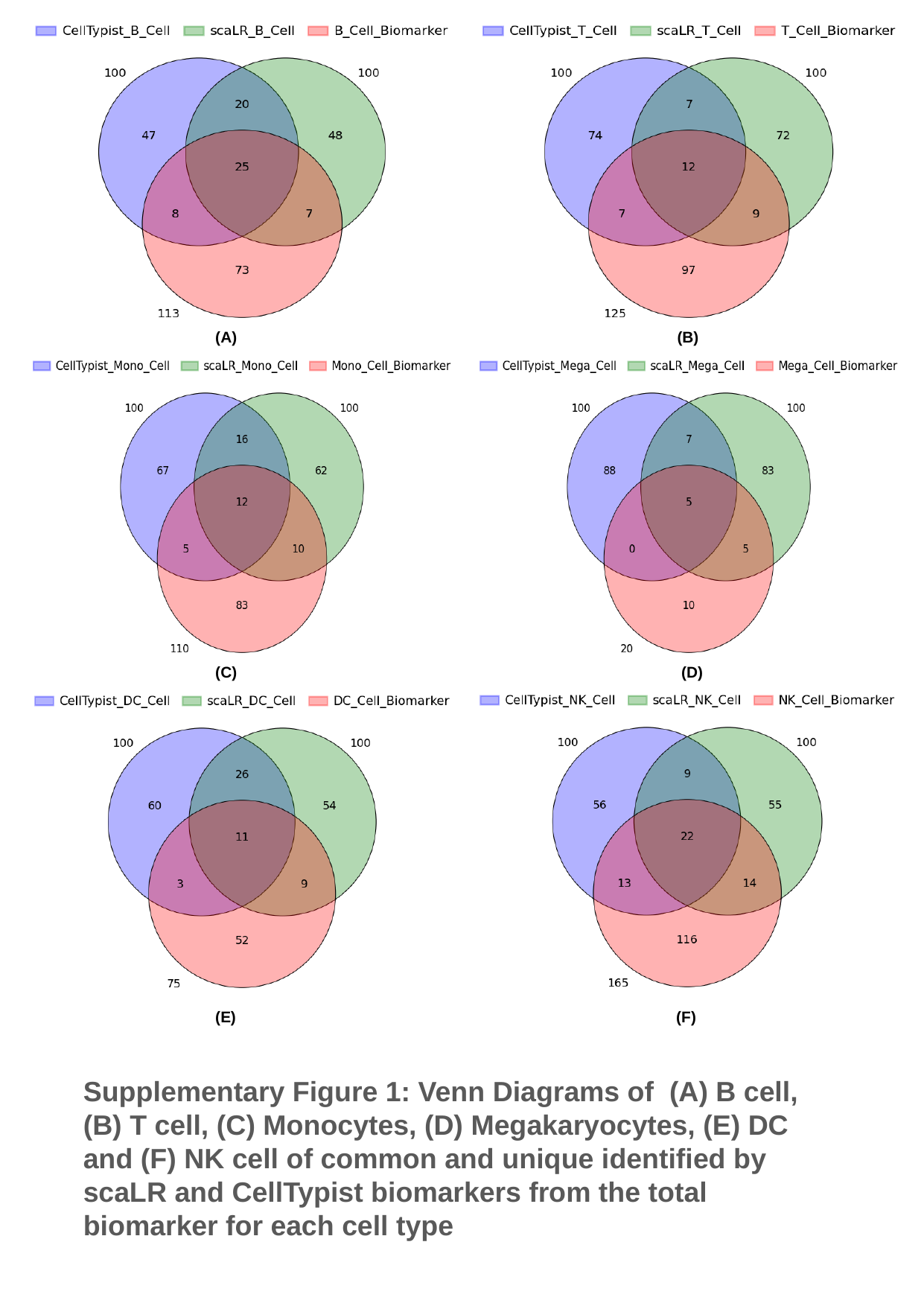

(A) (B)
(C) (D)
(E) (F)
Supplementary Figure 1: Venn Diagrams of (A) B cell, (B) T cell, (C) Monocytes, (D) Megakaryocytes, (E) DC and (F) NK cell of common and unique identified by scaLR and CellTypist biomarkers from the total biomarker for each cell type
